# Female vocalizations predict reproductive output in Brown-headed Cowbirds (*Molothrus ater*)

**DOI:** 10.1101/380642

**Authors:** Gregory M. Kohn

## Abstract

Pair bonds are often maintained through the reciprocal and coordinated exchange of communicative signals. The ability to recognize and appropriately respond to a partner’s signals will define a pair’s ability to reproduce. Individual variation in responsiveness, by shaping the formation and maintenance of strong pair bonds, will ultimately influence an individual’s reproductive output. Throughout the breeding period, female cowbirds (*Molothrus ater*) respond to male song displays using a vocalization known as the chatter. In this study, we investigated whether variation in chatters remained repeatable across years and predicted reproductive performance. A flock of cowbirds housed in a large aviary complex was observed during the spring of 2011 to 2012. We recorded courtship interactions, including singing behavior for males, and chatters and eggs laid by females. The rate with which females responded to song using chatters remained consistent across years, with some females predictably responding to more songs using chatters than others. During 2012, chattering predicted the number of eggs females laid and her paired status. Paired females were more likely to respond to songs with chatters, and there was a strong positive relationship between the number of eggs laid and the proportion of songs she responded to using chatters. Overall, these findings suggest that individual variation in female vocal responsiveness is an important contributing factor to cowbird reproductive success.

## INTRODUCTION

The ability to form and maintain pair bonds is a key factor in reproductive success (1–5). Successful pair bond maintenance requires pairs to coordinate activities and behavior to create strong, enduring, relationships. Within most vertebrate species, individuals possess social displays and vocalizations that attract the attention of, and coordinate activities with, potential or established mates (6). Individual differences in the use of such displays may create stronger social bonds with preferred mates, and ultimately increase reproductive output over time (7).

Increasingly, female displays and vocalizations are seen as critical factors shaping courtship and pair bonds in a wide array of species (8–11). During the breeding season, male cowbirds perform directed song displays at males and females. During song displays, males orient towards a neighboring individual and perform a song while spreading their wings and bowing (12). Cowbird courtship revolves around the female’s response to these song displays, and males modulate the intensity of their visual display in order to minimize female withdrawal (13). Females communicate their mate preferences using both visual (10) and acoustic (14) responses to male song displays. During the fall, males depend on these response displays for the development of their song, with females preferentially responding to, and reinforcing, high-quality song variants (10). Nevertheless, less is known about the factors shaping variation in such female responses, and how such variation predicts later reproductive outcomes for females.

Across many species females utilize vocalizations in response to male courtship displays (e.g., red winged blackbirds, *Agelaius phoeniceus* (15), grasshopper sparrows, *Ammodramus savannarum* (16), dunnocks, *Prunella modularis* (17), and duetting species (18, 19)). While female cowbirds do not sing, they possess an individually distinct chatter vocalization that is commonly used in response to a male’s song display (20). These response chatters often overlap or directly follow the end of a directed song display. In the wild, playbacks of chatters attract attention from both males and female cowbirds (21, 22); in the lab, females who are unselective in their chatters – by responding to playbacks of many different males’ song with chatters –are also less likely to maintain a pair bond (23). Females exposed to playbacks of songs followed by a playback of a response chatters also preferred those songs in contrast to females who were only exposed to playbacks of the song alone (14). These studies suggest that in cowbirds, as in many other species (24), the selective and reciprocal exchange of vocalizations across males and females plays a role in communicating mate preferences and maintaining pair bonds.

The aim of this study was to investigate whether consistent individual differences in the use of female vocalizations predict reproductive output in a semi-naturalistic flock setting. My first aim was to uncover if individual variation in female responsiveness remains repeatable, with some females consistently responding to more song display with chatters than others across two different breeding seasons. Across fall flock changes, female cowbirds exhibit consistent individual differences in the selectivity and frequency of their autumn social interactions (25) and use of affiliative head-down displays (26). Juvenile females who more frequently used affiliative “head-down” displays as juveniles during the fall were also more willing to respond to song using chatters during their first breeding season. This study will expand these findings to uncover if consistent individual differences in chatters are sustained across breeding seasons during adulthood.

My second aim was to uncover whether variation in the use of female vocalization reflects their reproductive output. Both strong pair bonds (7), and increased vocal responsiveness (27) can influence egg production in birds by stimulating and maintaining female reproductive physiology. As brood parasites, cowbirds do not raise their own young, and lay eggs in host species nests. Thus, the ability to place more eggs in more nests is crucial to gaining higher reproductive success. Cowbirds are also monogamous and maintain a single pair bond throughout the breeding season. I hypothesize that female cowbirds who consistently respond more to a higher proportion of song displays with chatters will be more likely to sustain a pair bond, and also exhibit higher rates of egg production than less responsive females.

## Methods

### Subjects

All birds were originally captured in Philadelphia County, Pennsylvania and Monroe County, Indiana and housed in aviaries in Monroe County, Indiana. All subjects were *Molothrus ater ater*. Previous studies have shown no differences in song or social behavior between the Philadelphia and Indiana populations (28). For this study we used 28 females including 21 adult (after second year by 2012) and 7 subadult (second year by 2012) females. We also used 28 males including 24 adult males and 4 subadult males. Birds ranged in age from 2 to 13 years old with an average age of 4.9 years. All birds had been used in previous studies, and were housed in large flocks prior to the beginning of this study. Each bird was marked with uniquely colored leg bands to allow for individual recognition. All birds were provided daily with a diet of vitamin-treated water (Aquavite Nutritional Research), red and white millet, canary seed and a modified Bronx Zoo diet for blackbirds.

### Aviaries

I used a single aviary complex that consisted of 4 subsections each with identical dimensions (9.1 x 21.4 x 3.4 meters), one small subsection (11 x 3 x 3.4 meters), and three indoor enclosures described in detail within Smith et al. (29). The large size of the aviary provides each cowbird with significant degrees of freedom to either engage or avoid interaction with conspecifics. Each large subsection of the aviary contained a covered feeding station and water bowls. Environmental conditions were similar throughout the entire aviary with shrubs, trees and grass that allowed individuals to both forage and hide. All birds were exposed to ambient climatic conditions, wild cowbirds, and the occasional sight of predators.

### Data collection

#### Behavioral observations

Throughout the study, a scan-sampling procedure was used to record behavioral observations; the entire flock was scanned and behaviors were recorded as they were observed (30). During scan sampling all behaviors were recorded using voice recognition technology described in detail by White, King & Duncan (31). When used in combination with voice recognition technology, scan-sampling can accurately acquire a more comprehensive dataset than focal sampling (32). All observations were conducted from 07:00-10:30 AM when cowbirds are most active, and were counterbalanced, so different observers took the same number of scan-sampling blocks in each aviary every day.

From June 9^th^ to July 8^th^ 2011 and from May 1^st^ to June 8^th^ 2012, we recorded courtship behavior, focusing on the vocal and approach behavior of both males and females. Throughout the study courtship behavior was recorded during 15-minute scan sampling blocks. For females, we recorded the number of songs each female received from males, and the number of female chatter vocalizations. Female chatter vocalizations were either response or undirected chatters. Response chatters occur when a female responds to a directed male song with chatter vocalization within a one second time window. Undirected chatter vocalizations occur when the females performs a chatter vocalization outside of singing contexts. For male courtship behavior, we recorded the number of female and male directed songs. Copulations were also recorded in order to assess female pair bonds (see below). During the pre-breeding season from March 18^th^ to April 23^rd^ in 2012 we also recorded approach behavior in separate 7-minute observation blocks. Here an approach was scored when one individual approached another individual with any part of its body within a radius of 30cm.

#### Egg Collection

From May 1^st^ to June 8^th^ we recorded the number of eggs each female laid. Six decoy nests were installed in each of the 4 large subsections of the aviary complex. Each nest was mounted on a forked perch attached to a backboard that contained a video camera, and was installed on posts or bushes within the aviary. All nests were supplied with yogurt-covered raisins as decoy eggs. A decoy egg was added every day to each nest until the nest contained three decoy eggs. Each day all nests were checked for the presence of cowbird eggs laid during the morning. After 8 days in one area each nest was moved to a different location within the aviary, nesting material was replaced, and was treated as a new nest starting with no eggs. All nests were video monitored to determine the identity of laying females by using Geovision software (Geovision Inc. 2008, 9235 Research Drive, Irvine, CA, USA) on Dell Vostro 230 computers running a 32-bit Windows 7 operating system. All work was conducted under ASAB/ABS guidelines and approved by the Institutional Care and Use Committee of Indiana University (08-018).

### Procedure

*Year 1: Spring 2011:* From June 9^th^ to July 8^th^ three observers collected a total of 240 observation blocks recording courtship behavior.

*Year 2: Spring 2012:* In the pre-breeding season from March 18^th^ to April 23^rd^, three observers collected a total of 40 blocks recording approach behavior and 164 blocks recording courtship behavior. During the breeding season from May 1^st^ to June 8^th^, three observers collected a total of 360 observational blocks recording courtship behavior. All decoy nest units were installed on May 1^st^ and used to record the number of eggs laid until the end of the breeding season on June 8^th^.

### Analysis

To document the repeatability of chatter across years, we used one-way intraclass correlation coefficients on the rate of each female’s chatters per block across 2011 and 2012. Intraclass correlation coefficients estimate the proportion of behavioral variance that is due to differences between individuals. To assess the rank ordered consistency in the individual tendency to chatter, we used Spearman’s correlations on the rate of response chatter across 2011 and 2012. All further analysis was conducted on the data recorded during spring 2012.

We considered a female to be paired if she received at least 100 songs and 70% of the songs she received came from a single male, with whom she exclusively copulated from 1 May to 8 June 2012. Furthermore, this female also had to be within the top two highest-ranking females sung to by the male. Thus, paired females maintained a selective relationship with a single male throughout the length of the breeding season, whereas unpaired females did not. We used Mann Whitney U-tests to look at the differences in the proportion of songs that a female responded to with a chatter, and the number of songs a female received between paired and unpaired females.

We used permutation-based linear models to investigate how variation in spring behavior predicted a female’s reproductive output. As social behavior often does not meet the assumption that errors are independent and normally distributed, permutation methods offer ideal alternatives to calculate probabilities of getting observed statistics after random reshuffling the data (33). For this study we used the lmp function in the lmPerm R package (34). I performed two models in this study: one model for all females, and another model restricted to paired females. Each model used an exact method to produce permutation probabilities and ran a minimum of 5000 permutations. As some explanatory factors were inter-correlated, we used variance inflation factors to assess the multicollinearity of main effects. A variance inflation factor greater than 10 is used to indicate potential multicollinearity, which makes model interpretation difficult (35). In none of our presented models did the VIFs for any main effects exceed 1.5. Post hoc analysis was conducted using Spearman’s correlations on continuous explanatory factors, and Wilcoxon rank sum test for categorical explanatory factors. Confidence intervals for Spearman’s coefficients were calculated using resampling techniques.

For both models, the dependent factor was the number of eggs that each female laid. For the all-female model, the explanatory factors included main effects of the total rate of songs received, paired status, the number of approaches initiated during the prebreeding season, proportion chatter (number of response chatters/ total number of songs), and their age class (sub-adult and adult) and the number of undirected chatters. The paired-female model was restricted to only females in a pair bond, and focused on how interactions in pairs predicted female reproductive output. The explanatory factors for the paired model were the rate of songs received from their paired male, the proportion chatter in response to their paired male, the female’s age class (sub-adult and adult), whether they were paired with the same or different male across years (same pair, different pair), and the number of undirected chatters.

## Results

### Repeatability of chatters across years

Across years, females were predictable in their propensity to respond to song displays using chatters. In 2011, we observed a total of 4,152 chatters including 1,272 response chatters (*Median per individual* = 28.5) and 2,880 undirected chatters (*Median per individual* = 28). During the breeding season in 2012, we observed a total of 6,830 chatters, including 2,339 response chatters (*Median per individual* = 27), and 4,491 undirected chatters (*Median per individual* = 36). For all females, individual variation in the rate of response chatters was repeatable across both years (*ICC* = 0.50, *p* < 0.0001, 95 % CI = 0.17–0.73). Females also showed significant rank-ordered consistency in the rate of response chatter in relation to other females across years (Spearman’s rank correlation: *rho* = 0.43, *N* = 28, *p* = 0.03, 95% CI = 0.06 – 0.73). Within both spring 2011 and 2012, females who performed the most undirected chatters also performed the most response chatters (2011: *rho* = 0.90, *N* = 28, *p* < 0.0001, 95% CI = 0.80 – 0.94, 2012: *rho* = 0.93, *N* = 28, *p* < 0.0001, 95% CI = 0.85 – 0.97).

### Chatters and pair bonds

Response chatters were used very selectively, and were primarily directed towards a single male across the breeding season. From 1 May to 8 June in 2012, we recorded 5,091 songs sung to females, with a median of 177.5 songs per female. For each female, we rank ordered the number of response chatters to each male and calculated the proportion of response chatters in response to each male’s songs. The top male accounted for the majority of the female’s response chatters (*Median proportion of response chatter to top male* = 0.90), and in paired females the top male was always the female’s partner. While paired females received more songs than unpaired females (*Median Paired Females* = 242, *Median Unpaired females* = 62, Mann-Whitney *U* test: *U* = 44.5, *N_1_* = 14, *N_2_* = 14, *p* = 0.0003), they were also more likely to respond to a higher proportion of songs with response chatters (*Median Paired Females* = 0.60, *Median Unpaired females* = 0.05, *U* = 14, *N_1_* = 14, *N_2_* = 14, *p* = 0.0001, Fig 1).

**Figure 1.**
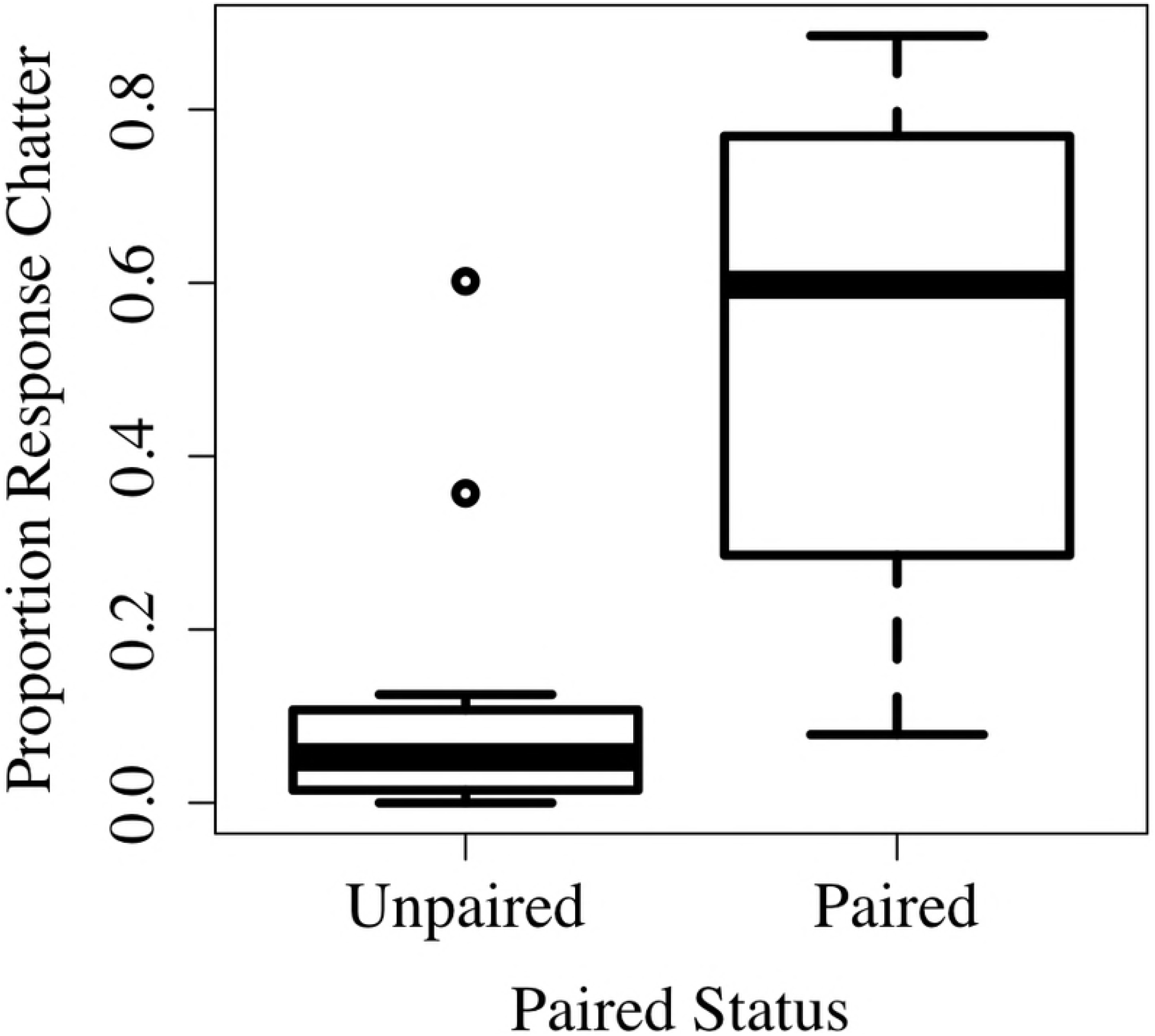
The proportion of response chatter vocalizations based on based on an individual’s paired status. Boxes represent interquartile ranges with the median in the middle represented by a bold line; whiskers represent the range of the highest and lowest values that are within a range of 1.5 times the interquartile range; dots indicate data points that are outside this range.

### Egg output All-Female Model

During the breeding season, females who laid more eggs responded to a higher proportion of songs with a response chatter. We identified the laying female for 93 eggs (*Mean eggs laid* = 3.32). Our model (Table1) explained 74% of the variance in eggs laid (*R*^2^= 0.74, *F_(7,20)_* = 8.12, *p* = 0.0001). The proportion of male song displays followed by a chatter was the only significant predictor of the number of eggs an individual laid (Table 1). Post hoc correlations revealed a significant positive relationship between the numbers of eggs an individual laid and proportion chatter (*rho* = 0.77, *N* = 28, *p* < 0.0001, 95% CI = 0.54 – 0.92, Fig 2). Additional analysis also showed that the rate of response chatters before the breeding season (before females were actively laying eggs), from 18 March to 23 April, was also positively correlated with the later number of eggs an individual laid (*rho* = 0.68, *N* = 28, *p* < 0.002, 95% CI = 0.43 – 0.84).

**Fig 2:**
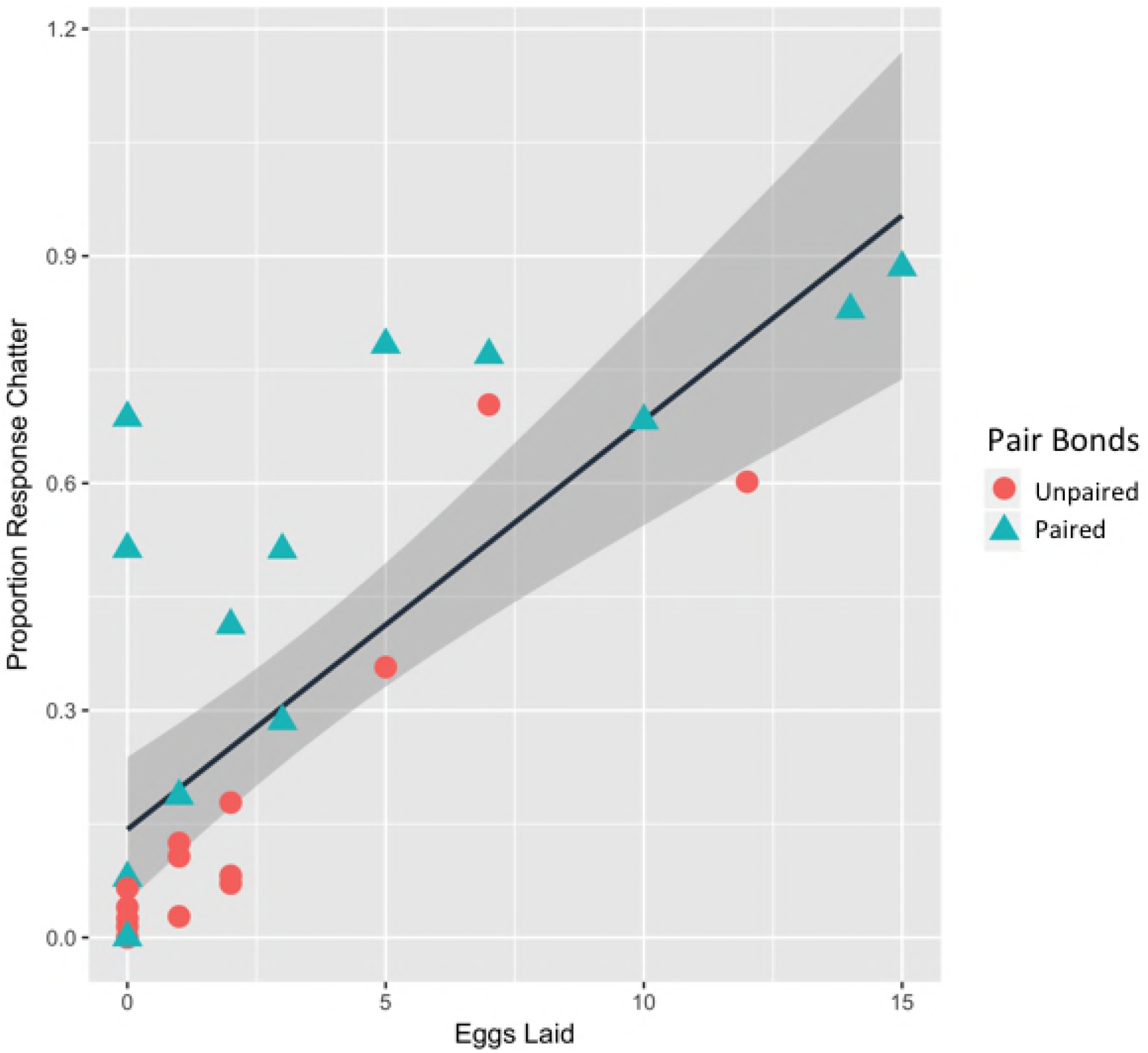
Scatterplots for the proportion of response chatters and the number of eggs laid for all females. Females who formed a pair bond during 2012 season are shown as a triangle, and females who did not maintain a pair bond are shown as a circle. Line represents the permuted linear regression with surrounding 95% confidence intervals.

**Table 1:** Results of the permutation-based linear models for eggs laid during the breeding season of 2017. Table represents the model for (A) all-females and (B) paired-females.

| A. All- Female Model | Coefficients | P value | A. Paired- Female Model | Coefficients | P value |
| --- | --- | --- | --- | --- | --- |
| **Songs Received** | -0.01 | p = 0.08 | Paired male song | -0.02 | p = 0.16 |
| **Approach** | 0.005 | p = 0.41 | Approach | 0.002 | p = 0.86 |
| **Proportion chatter** | 14.65 | p < 0.00001*** | Proportion paired chatter | 12.47 | p = 0.03* |
| **Age class** | 0.95 | p = 0.50 | Age Class | 4.97 | p = 0.11 |
| **Undirected chatter** | 0.58 | p = 0.69 | Undirected Chatter | 2.27 | p = 0.24 |
| **Pair bond** | 1.28 | p = 0.38 | Stable/ Switched pair bonds | 2.69 | p = 0.20 |

We identified 72 eggs from adult females (*Mean* = 3.42) and 21 eggs from subadult females (*Mean* = 3). Age did not significantly influence the number of eggs produced. There was no significant difference in the number of eggs produced by subadults in contrast to adults (*Median Adult* = 2.00, *Median Subadult* = 0.05, *N_1_* = 21, *N_2_* = 7, *U* = 88, *p* = 0.45). While paired status did not reach significance in our model, post hoc analysis revealed that paired females produced more eggs than unpaired females (*Median Paired* = 3.00, *Median Unpaired* = 0.05, *U* = 145, *N_1_* = 14, *N_2_* = 14, *P* = 0.03).

### Egg output Paired-Female model

Our paired-female model explained 78% of the variance in egg laying (*R*^2^= 0.78, *F_(5,8)_* = 5.683, *p* = 0.016) and had only one significant predictor, the proportion of songs followed by a response chatter (Table 1). None of the other variables were significant predictors of the number of eggs a female laid (Table 1). Within paired individuals, the proportion of response chatters was significantly correlated with the number of eggs laid (*rho* = 0.72, *p* = 0.004, 95% CI = 0.33 – 0.92, Fig 2), but neither the number of undirected chatters (*rho* = 0.42, *p* = 0.13, 95% CI = −0.10 – 0.83), nor the number of songs they received from their paired male (*rho* = −0.03, *p* = 0.92, 95% CI = −0.55 – 0.48).

In order to look at the factors predicting variation in response chatters I conducted an additional permuation based linear model. The dependent variable in this model was the proportion of response chatters to her paired males songs. The explanatory factors were age, songs received from paired males, and if the female maintained a stable pair bond across breeding seasons. This model was not significant (*R*^2^= 0.22, *F_(3,10)_* = 0.93, *p* = 0.46). The number of songs a female received from her paired male was not significantly correlated with proportion of response chatters (*rho* = 0.37, *N* = 14, *p* = 0.19, 95% CI = −0.06 – 0.68). There was also no significant differences in both the proportion of response chatters (W = 28, *p* = 0.662), the number of eggs laid (W = 33.5, *p* = 0.24) between females who were paired with the same male across both breeding seasons, and females who changed males.

## Discussion

I investigated the association between individual differences in courtship behavior and reproductive performance in female brown-headed cowbirds. Female cowbirds exhibited consistent individual differences in their responsiveness to male song, with some females being more likely to respond to male song displays using chatters than others. As vocal stimuli are important for attracting potential partners (36), shaping reproductive physiology (37, 38), and maintaining pair bonds (15, 39), consistency in vocal responsiveness may reliably construct the social relationships needed for increased reproductive output. In accordance with this, I discovered that the proportion of song displays a female responded to with chatters was greater in paired females, and predicted the number of eggs she produced. In paired females, I also found that the proportion of response chatters to their paired male’s song display was the only significant predictor of the number of eggs she laid.

Paired females responded to a higher proportion of songs with chatters than unpaired females. This suggests that the maintenance of pair bonds is associated with the reciprocal exchange of vocal displays from both male and female cowbirds. While recognition of female courtship displays is becoming more widespread (40, 41), little is currently known about how these displays shape their relationship with males. Pervious studies have shown have shown that increased attention, coordination, and synchrony within pairs has multiple benefits, such as increasing vigilance, lowering the energetic demands of foraging and parental care, and more effective mate guarding (42–44). In alpine accentors (*Prunella collaris*) females use complex songs to attract mates (36), and the calls of female whitethroats (*Sylvia communis*) both attract males and shape their courtship behavior (45). In many mammals such as brown rats, *Rattus norvegicus*, (46), grey mouse lemurs, *Microcebus murinus*, (47), and Barbary Macaques, *Macaca sylvanus*, (48), female vocalizations often reflect reproductive status, and are used to attract males. In the field, playbacks of cowbird chatters often attract males to the location of a speaker (22), and males will often follow and peruse females who responded to their song with a chatter (Kohn, personal observation). By possessing a signal that reflects their reproductive status, female cowbirds who are more vocally responsive will be better able to attract preferred male attention and drive pair coordination across the breeding season.

Variation in signals used to attract and coordinate activities within pairs can have cascading influences on later survival and fitness. I found that a female’s vocal response to male song displays was the strongest predictor of her reproductive output, with more vocally responsive females laying more eggs than less responsive females. Similar findings have been observed in red-winged blackbirds, where females who had a successful nest were more likely to answer male songs with a chit vocalization (39). In many species, the reciprocal displays between members of a pair can also shape reproductive physiology (27, 49). For instance, in ring doves (*Streptopelia risoria*), the presence of a preferred male song stimulates the females to use ‘coo’ vocalizations (50). In turn, the coo vocalizations themselves stimulate ovarian development (27, 49), which may result in increased egg production. Thus, the contingent displays females use in response to their partners may be an important, albeit under-recognized, component in shaping a pair’s reproductive success.

Currently, the direction of effects between increased reproductive output and coordinated displays between cowbird pairs is unknown. However, females begin responding to male song with chatters prior to the egg laying period, and response chatter rates during this pre-laying period are correlated with egg output the same year. Thus, a female’s own courtship behavior might play a role in providing the necessary stimulation for increased reproductive output. While the mechanisms underlying the relationship between vocal responsiveness, pair bonds, and egg production need further investigation, my results demonstrate that repeated use of response chatters is predictive of increased reproductive output in female cowbirds.

In cowbirds, female responses to male vocalizations are commonly used to assess the quality and attractiveness of male signals (10). Females use their response chatter selectively, almost exclusively in response to their paired males. As females exclusively copulated with their paired males, response chatters may be a reliable signal of female preferences, and used to reinforce pair bonds. Chatters are also individually distinct (20), and their selective use may facilitate the individual identification needed to sustain a monogamous pair bond (15, 51). Female cowbirds with lesions to their HVC area are not selective in their response chatters, and chatter in response to nearly all song playbacks, regardless of their quality (23). These lesioned females are also unable to sustain a pair bond, and are courted by a larger number of males than other females. I found that females who retained the same pair-bonded males across two different breeding seasons showed no significant differences in vocal responsiveness or egg production when compared to females who changed paired males. The number of songs a female received from males did not reflect the proportion of response chatters to his songs, and further analysis also showed that the number of response chatters a male received across breeding seasons was not correlated or repeatable (Sup 1). While the correlational nature of this study does not allow us to directly test how differences in male quality or song can influence female vocal responses, our result suggest that variation in the use of chatters represents different behavioral strategies that females use when engaging and forming pair bonds with preferred males.

This paper adds to the increasing number of studies showing the importance of female vocalizations in constructing and reinforcing avian pair bonds (52, 53), and further suggests that female vocalizations contributes to their reproductive success. Consistent individual differences in cowbird social behavior can predict an individual’s reproductive performance across long timescales (54). Juvenile female cowbirds who initiate more affiliative head-down displays during autumn are more likely to engage males with chatters and form a pair bond during their first breeding season (26). Here we show that such variation in female vocal responses is maintained into adulthood, remains associated with pair-bond status, and predicts reproductive output. In cowbirds, social experiences are critical in the development female mate preferences (55, 56), and may also shape behavioral differences in how females interact with preferred males (26). Further research will explore how the early social environment shapes the development of individual differences in chatter vocalizations among females, and the causal mechanisms linking chatter vocalizations, pair bonds, and increased reproductive output.

## ACKNOWLEDGEMENTS

All work was conducted under ABS guidelines and approved by the Institutional Care and Use Committee of Indiana University (08-018). I would like to thank Meredith West and Andrew King for their support and advice during this project.

